# TEAD4/YAP1/WWTR1 prevent the premature onset of pluripotency prior to the 16-cell stage

**DOI:** 10.1101/663005

**Authors:** Tristan Frum, Jennifer Watts, Amy Ralston

## Abstract

In the mouse embryo, pluripotent cells arise inside the embryo around the 16-cell stage. During these early stages, *Sox2* is the only gene whose expression is known to be induced specifically within inside cells as they are established. To understand how pluripotent cells are created, we investigated the mechanisms regulating the initial activation of *Sox2* expression. Surprisingly, *Sox2* expression initiated normally in the absence of both *Nanog* and *Oct4*, highlighting differences between embryo and stem cell models of pluripotency. However, we observed precocious, ectopic expression of *Sox2* prior to the 16-cell stage in the absence of *Yap1, Wwtr1*, and *Tead4*. Interestingly, the repression of premature *Sox2* expression was sensitive to LATS1/2 activity, even though it normally does not limit TEAD4/YAP1/WWTR1 activity during these early stages. Finally, we present evidence for direct transcriptional repression of *Sox2* by YAP1/WWTR1/TEAD4. Taken together, our observations reveal that, while embryos are initially competent to express *Sox2* as early as the 4-cell stage, transcriptional repression prevents the premature expression of *Sox2*, thereby restricting the pluripotency program to the stage when inside cells are first created.

## Introduction

Pluripotency describes the developmental potential to produce all adult cell types. However, in mammals, the establishment of pluripotency takes place in the context of lineage decisions that separate pluripotent cells of the fetus from cells that give rise to extraembryonic tissues such as the placenta. Thus, in mammals, the onset of pluripotency is initially delayed as the blastomeres transition from totipotency to adopt the more specialized pluripotent and extraembryonic states.

The mouse embryo has provided an invaluable tool to understand the molecular mechanisms that initially create pluripotent cells, which are also the progenitors of embryonic stem cells. While much progress has been made in understanding how pluripotency is maintained once pluripotent cells are established, the mechanisms driving the initial establishment of pluripotency remain relatively obscure.

In the mouse embryo, pluripotent cells emerge from cells positioned inside the embryo, which occurs around the 16-cell stage, and continues as the inside cells form the inner cell mass of the blastocyst. The inner cell mass will go on to differentiate into either pluripotent epiblast or non-pluripotent primitive endoderm. As the epiblast matures, it gradually acquires a more embryonic stem cell-like transcriptional signature (Boroviak et al., 2014; Boroviak et al., 2015).

While studies in mammalian embryos and embryonic stem cells have developed an extensive catalog of transcription factors that promote pluripotency, the only pluripotency-promoting transcription factor known to distinguish inside cells as they form at the 16-cell stage is *Sox2* (Guo et al., 2010; Wicklow et al., 2014). Therefore, understanding how *Sox2* expression is regulated at the 16-cell stage can provide insight into how pluripotency is first established.

Here, we use genetic approaches to test mechanistic models of the initial activation *Sox2* expression. We investigate the contribution, at the 16-cell stage and prior, of factors and pathways that are known to regulate expression of *Sox2* at later preimplantation stages and in embryonic stem cells. We show that embryos are competent to express *Sox2* as early as the four-cell stage, although they normally do not do so. Finally, we uncover the molecular mechanisms that ensure that *Sox2* expression remains repressed until the developmentally appropriate stage.

## Results and Discussion

### The initiation of *Sox2* expression is *Nanog*- and *Oct4*-independent

To identify mechanisms contributing to the onset of *Sox2* expression in the embryo, we first focused on the role of transcription factors that are required for Sox2 expression in embryonic stem cells. The core pluripotency genes *Nanog* and *Oct4* (*Pou5f1*) are required for *Sox2* expression in embryonic stem cells (Chambers et al., 2003; Mitsui et al., 2003; Niwa et al., 2005) and are expressed at the 8-cell stage (Dietrich and Hiiragi, 2007; Palmieri et al., 1994; Rosner et al., 1990; Strumpf et al., 2005), prior to the onset of *Sox2* expression at the 16-cell stage, suggesting that NANOG and OCT4 could activate the initial expression of *Sox2*.

We previously showed that the initiation of *Sox2* expression is *Oct4*-independent, as embryos lacking *Oct4* have normal levels of *Sox2* expression at E3.5 (Frum et al., 2013). We therefore hypothesized that *Nanog* and *Oct4* could act redundantly to initiate *Sox2* expression. To test this hypothesis, we bred mice carrying the null allele *Nanog-GFP* (Maherali et al., 2007) with mice carrying an *Oct4* null allele (Kehler et al., 2004) to generate *Nanog;Oct4* null embryos (Fig. S1A). In wild-type embryos, *Sox2* is first detected in inside cells at the 16-cell stage, with increasing robustness in inside cells of the 32-cell stage embryo (Guo et al., 2010; Wicklow et al., 2014). In *Nanog;Oct4* null embryos, SOX2 was detectable at the 16-cell (E3.0) and 32-cell (E3.25) stages (Fig. 1A-B). We observed no differences in the proportions of SOX2-expressing cells at the 16- and 32-cell stages between non-mutant embryos and embryos lacking *Nanog, Oct4*, or both (Fig. S1B,C). These observations indicate that *Nanog* and *Oct4* do not regulate initial *Sox2* expression.

**Figure 1.**
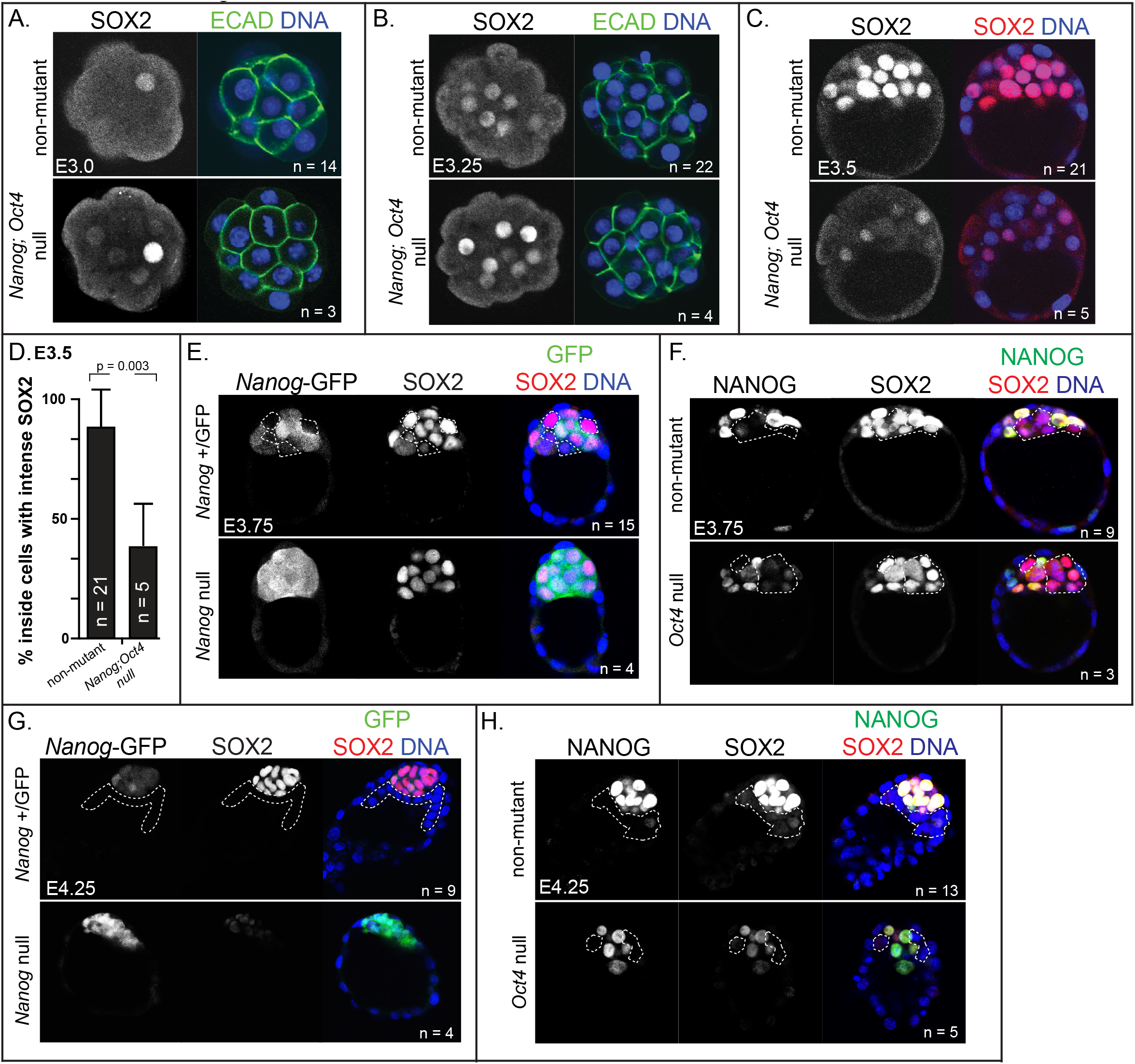
*Nanog* and *Oct4* are required for the maintenance, but not the initiation of *Sox2*. (A) Immunostaining for SOX2, E-Cadherin (ECAD), and DNA in non-mutant and *Nanog;Oct4* null embryos at the 16-cell stage (E3.0). (B) SOX2, ECAD, and DNA in non-mutant and *Nanog;Oct4* null embryos at the 32-cell stage (E3.25). (C) SOX2 in non-mutant and *Nanog;Oct4* null embryos at E3.5. (D) Quantification of the percentage of inside cells, across all embryos, with intense SOX2 staining in non-mutant and *Nanog;Oct4* null embryos at E3.5. Columns = mean, error bars = standard deviation, p = Student’s t-test. (E) NANOG-GFP, SOX2, and DNA in non-mutant and *Nanog* null embryos at E3.75. (F) NANOG, SOX2, and DNA in non-mutant and *Oct4* null embryos at E3.75. (G) NANOG-GFP, SOX2, and DNA in non-mutant and *Nanog* null embryos at E4.25. (H) NANOG, SOX2, and DNA in non-mutant and *Oct4* null embryos at E4.25. For all panels, n = number of embryos examined. Dashed white lines demarcate non-epiblast/presumptive primitive endoderm cells.

### *Nanog* and *Oct4* are individually required to maintain *Sox2* expression

To investigate a role for *Nanog* and *Oct4* in maintaining expression of *Sox2*, we evaluated double null embryos at a later time point. By E3.5, SOX2 appeared weak or undetectable in most cells of *Nanog;Oct4* null embryos (Fig. 1C). Moreover, the proportion of cells expressing the wild type level of SOX2 was significantly lower in *Nanog;Oct4* null embryos (Fig. 1D), but not in embryos lacking *Nanog* or *Oct4* only (Fig. S1D). We therefore conclude that *Nanog* and *Oct4* work together to maintain *Sox2* expression and can compensate for the loss of one another up to at least E3.5.

To evaluate whether *Nanog* and *Oct4* cooperatively maintain *Sox2* expression at later preimplantation stages, we examined SOX2 expression in embryos lacking either *Nanog* or *Oct4* at later developmental stages. At E3.75, SOX2 levels were indistinguishable between non-mutant, *Nanog* null and *Oct4* null embryos (Fig 1E,F). In fact, the only difference between non-mutant, *Oct4* null and *Nanog* null embryos at this timepoint was the previously reported failure of *Nanog* null embryos to undergo primitive endoderm differentiation by downregulating *Nanog* expression in a subset of inner mass cells (Frankenberg et al., 2011; Messerschmidt and Kemler, 2010).

By contrast, both *Nanog* null and *Oct4* null embryos exhibited defects in SOX2 by E4.25. *Nanog* null embryos exhibited the more severe SOX2 expression phenotype, with almost undetectable SOX2 (Fig. 1G). *Oct4* null embryos exhibited a less severe phenotype, with reduced, but detectable SOX2 (Fig. 1H). These observations indicate that, while the initial phase of *Sox2* expression is independent of *Nanog* and *Oct4*, this is followed by a period during which *Nanog* and *Oct4* act redundantly to maintain *Sox2* expression, which then gives way to a phase during which *Nanog* and *Oct4* are individually required to achieve maximal *Sox2* expression.

### TEAD4/WWTR1/YAP1 regulate the onset of *Sox2* expression

Having observed that the initiation of *Sox2* expression is *Nanog*- and *Oct4*-independent, we next examined the role of other factors in regulating initial *Sox2* expression. TEAD4 and its co-factors WWTR1 and YAP1 repress *Sox2* expression in outside cells starting around the 16-cell stage (Frum et al., 2018; Wicklow et al., 2014). However, YAP1 is detected within nuclei as early as the 4-cell stage (Nishioka et al., 2009), suggesting that the complex is active prior to the 16-cell stage. Recent studies have highlighted the roles and regulation of TEAD4/WWTR1/YAP1 in promoting outside cell maturation to trophectoderm during blastocyst formation (Anani et al., 2014; Cao et al., 2015; Cockburn et al., 2013; Hirate et al., 2013; Kono et al., 2014; Leung and Zernicka-Goetz, 2013; Lorthongpanich et al., 2013; Nishioka et al., 2009; Nishioka et al., 2008; Posfai et al., 2017; Rayon et al., 2014; Shi et al., 2017; Yagi et al., 2007; Yu et al., 2016). Yet, the developmental requirement for TEAD4/WWTR1/YAP1 prior to the 16-cell stage has not been investigated. We therefore hypothesized that TEAD4/WWTR1/YAP1 repress *Sox2* expression prior to the 16-cell stage.

To test this hypothesis, we examined SOX2 in embryos lacking *Tead4*. Consistent with our hypothesis, *Tead4* null embryos exhibited precocious SOX2 at the 8-cell stage (Fig. 2A,C). Notably, this phenotype that was not exacerbated by elimination of maternal *Tead4* (Fig. S2A and Fig. 2A,C), consistent with the absence of detectable *Tead4* in oocytes (Yagi et al., 2007). By contrast, deletion of maternal *Wwtr1* and *Yap1* (Fig. S2B) led to precocious SOX2 at the 8-cell stage (Fig. 2B,D). The presence of wild-type, paternal alleles of *Wwtr1* and/or *Yap1* did not rescue SOX2 in the maternally null embryos (Fig. 2B, D). Therefore, maternally provided WWTR1/YAP1 and zygotically expressed TEAD4 repress *Sox2* expression at the 8-cell stage.

**Figure 2.**
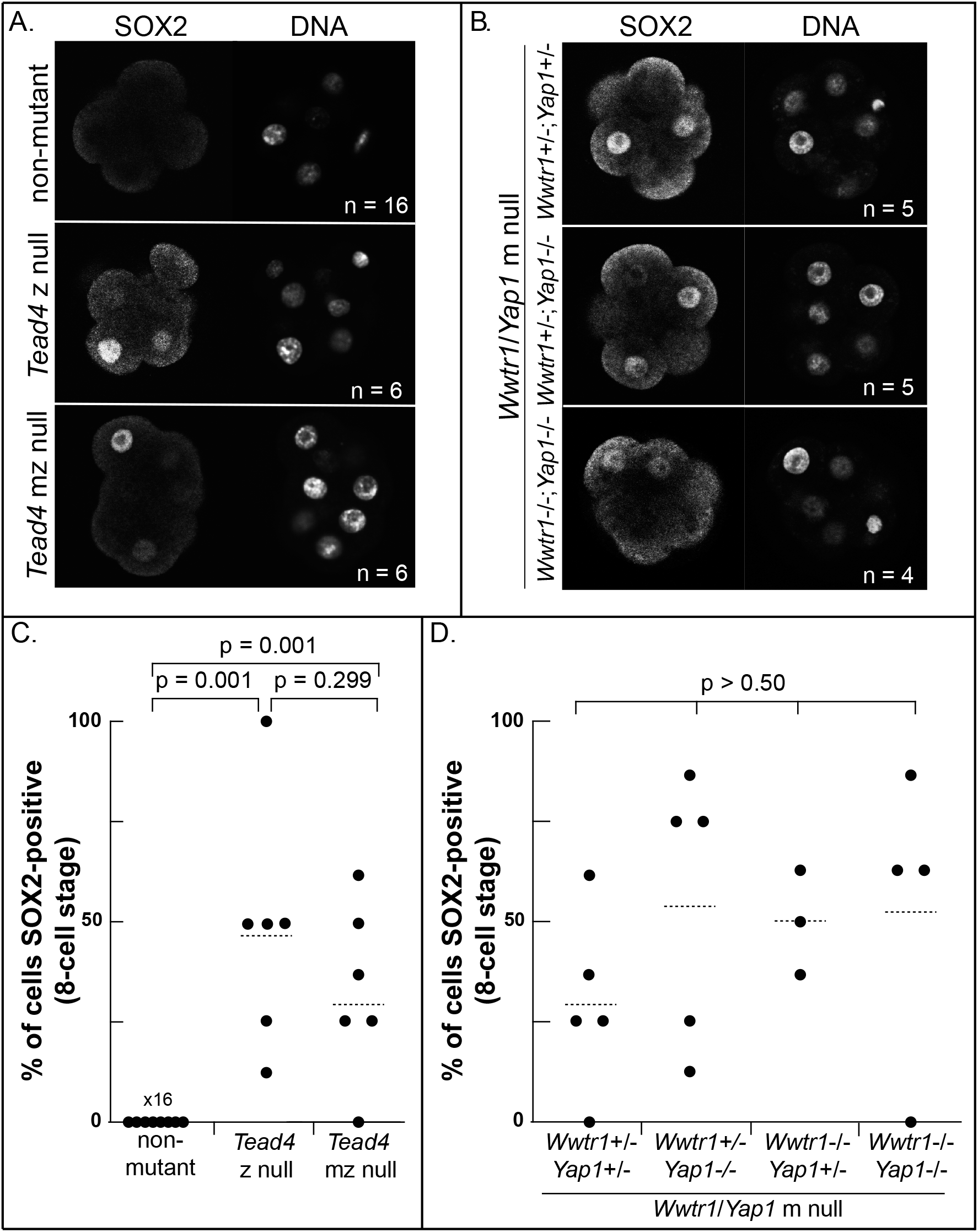
TEAD4/WWTR1/YAP1 repress precocious *Sox2* expression at the 8-cell stage. (A) Immunostaining for SOX2 in non-mutant, *Tead4* zygotic (z) null and *Tead4* maternal-zygotic (mz) null embryos at the 8-cell stage. (B) SOX2 in embryos lacking m *Wwtr1* and *Yap1* at the 8-cell stage, with indicated zygotic genotypes. (C) The percentage of SOX2-positive cells per embryo (dots) in non-mutant, *Tead4* z null and *Tead4* mz null embryos. (D) The percentage of SOX2-positive cells per embryo (dots) in *Wwtr1/Yap1* m null embryos with indicated zygotic genotype. Dashed line = mean, p = one-way ANOVA with Tukey post-hoc test, n = number of embryos examined.

We next evaluated SOX2 in embryos lacking maternal *Wwtr1;Yap1* at the 4-cell stage. We observed that 4-cell embryos lacking maternal *Wwtr1* and *Yap1* occasionally exhibited weak ectopic SOX2 (Fig. S2D,E). However, SOX2 was never detected in 4-cell embryos lacking *Tead4* (Fig. S2C). These observations suggest that *Wwtr1* and *Yap1* partner with other factors to regulate the onset of *Sox2* expression at the 4-cell stage and point to a requirement for other TEAD proteins that are expressed at the 4-cell stage (Nishioka et al., 2008).

The premature onset of *Sox2* expression in embryos lacking *Tead4* or *Wwtr1* and *Yap1* demonstrates that preimplantation mouse embryos are capable of expressing markers of inside cell identity as early as the 4-cell stage and reveals an earlier than expected role for TEAD4/WWTR1/YAP1 in repressing the expression of *Sox2* until the formation of inside cells, thus permitting the establishment of discrete trophectoderm and inner cell mass gene expression. Furthermore, the appearance of *Sox2* expression prior to the formation of inside cells argues that no cues specific to inside-position are required for *Sox2* expression beyond regulated TEAD4/WWTR1/YAP1 activity. Rather, our results suggest that the mechanism regulating the onset of Sox2 expression is that constitutive repression of *Sox2* by TEAD4/WWTR1/YAP1 is relieved once cells are positioned inside the embryo at the 16-cell stage.

### Repression of *Sox2* at the 4- and 8-cell stage is sensitive to LATS2 kinase

In many contexts, TEAD4/WWTR1/YAP1 activity is repressed by the HIPPO pathway LATS1/2 kinases, which repress nuclear localization of WWTR1/YAP1 (Nishioka et al., 2009; Zhao et al., 2010; Zhao et al., 2007). To evaluate the role of HIPPO signaling in regulating initial *Sox2* expression, we therefore examined whether *Sox2* expression is LATS1/2-sensitive prior to the 16-cell stage.

We injected mRNA encoding *Lats2* into both blastomeres of 2-cell stage embryos, which is sufficient to inactivate the TEAD4/WWTR1/YAP1 complex during blastocyst formation (Nishioka et al., 2009; Wicklow et al., 2014), and then evaluated SOX2 at 4- and 8-cell stages (Fig. 3A). As anticipated, *Lats2* mRNA injection, but not injection of *Green Fluorescent Protein* mRNA, greatly reduced YAP1 nuclear localization at 4- and 8-cell stages (Fig. 3B,C). In addition, we observed precocious SOX2 in embryos overexpressing *Lats2* (Fig 3B,C). Therefore, LATS kinases can prevent TEAD4/WWTR1/YAP1 from repressing expression of *Sox2* prior to the 16-cell stage, but LATS kinases must not normally do so, since *Sox2* is usually repressed during early development.

**Figure 3.**
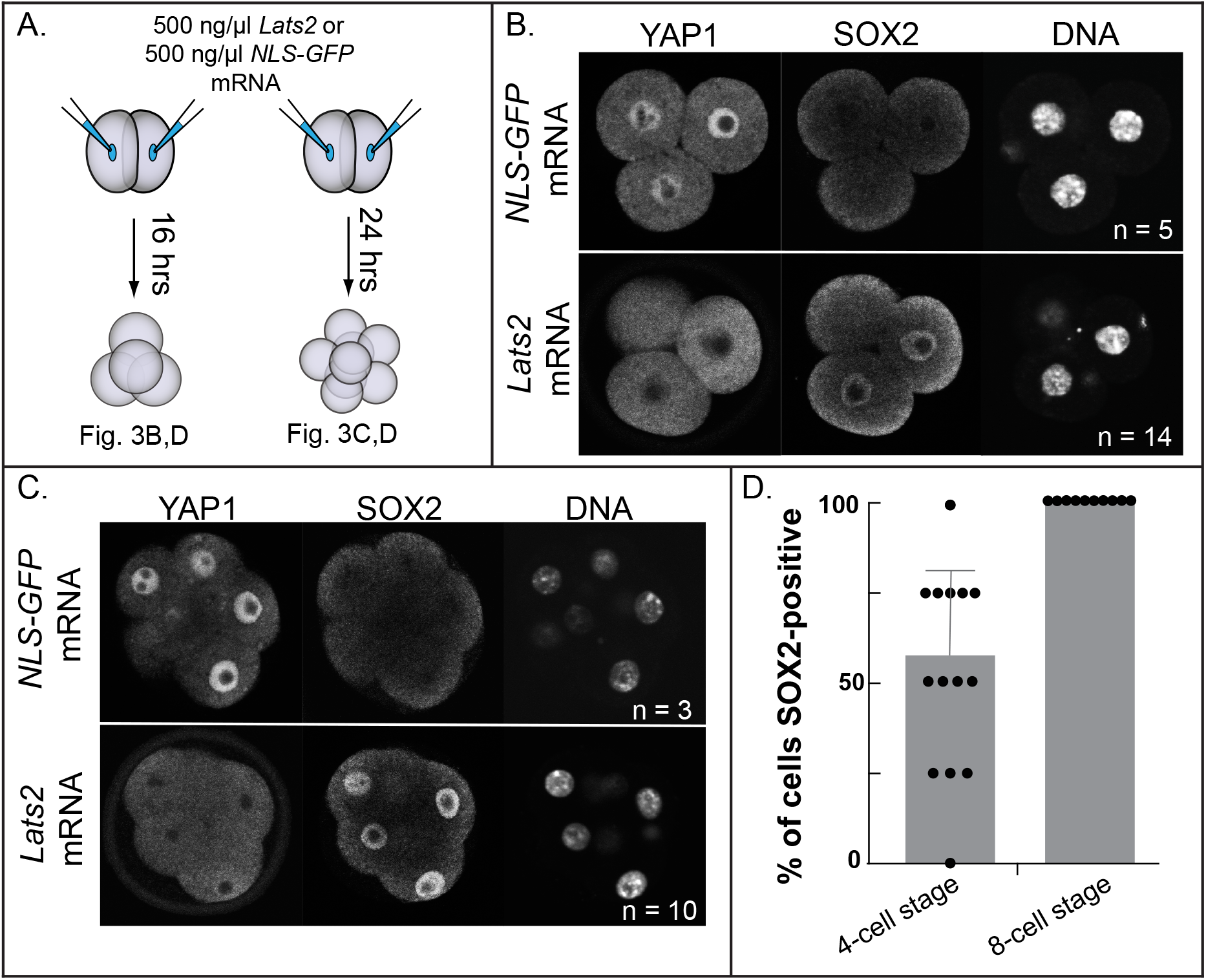
YAP1 localization and *Sox2* expression are sensitive to LATS2 kinase. (A) Experimental approach: both blastomeres of 2-cell stage embryos were injected with either 500 ng/μl *NLS-GFP* mRNA, which encodes GFP bearing a nuclear localization sequence (NLS), or 500 ng/μl *Lats2* mRNA, and were then cultured to the 4- or 8-cell stages. (B) YAP1 and SOX2 immunostaining in 4-cell stage embryos injected with *NLS-GFP* mRNA or *Lats2* mRNA. (C) YAP1 and SOX2 in 8-cell stage embryos injected with *NLS-GFP* mRNA or *Lats2* mRNA. (D) Dot-plot of the percentage of SOX2-positive cells per embryo (dots) at the indicated stages. The means and standard deviations are represented by columns and error bars, while n = number of embryos examined.

We observed in published RNA-seq data sets that *Lats1* and *Lats2* are expressed in 4-cell and 8-cell stage embryos (Tang et al., 2010; Wu et al., 2016), opening an interesting future direction of discovering how TEAD4/WWTR1/YAP1 escape inhibition by LATS kinases prior to the 16-cell stage, which is not currently understood.

### TEAD4/WWTR1/YAP1 may repress *Sox2* expression through a direct mechanism

While the TEAD4/WWTR1/YAP1 complex is widely recognized as a transcriptional activator, it has more recently been shown to act as a transcriptional repressor (Beyer et al., 2013; Kim et al., 2015). Therefore, we considered two mechanisms by which TEAD4/WWTR1/YAP1 could repress *Sox2* expression (Fig. 4A): an indirect model, in which TEAD4/WWTR1/YAP1 promote transcription of a *Sox2* repressor, and a direct model, in which TEAD4/WWTR1/YAP1 themselves act as the *Sox2* repressor.

**Figure 4.**
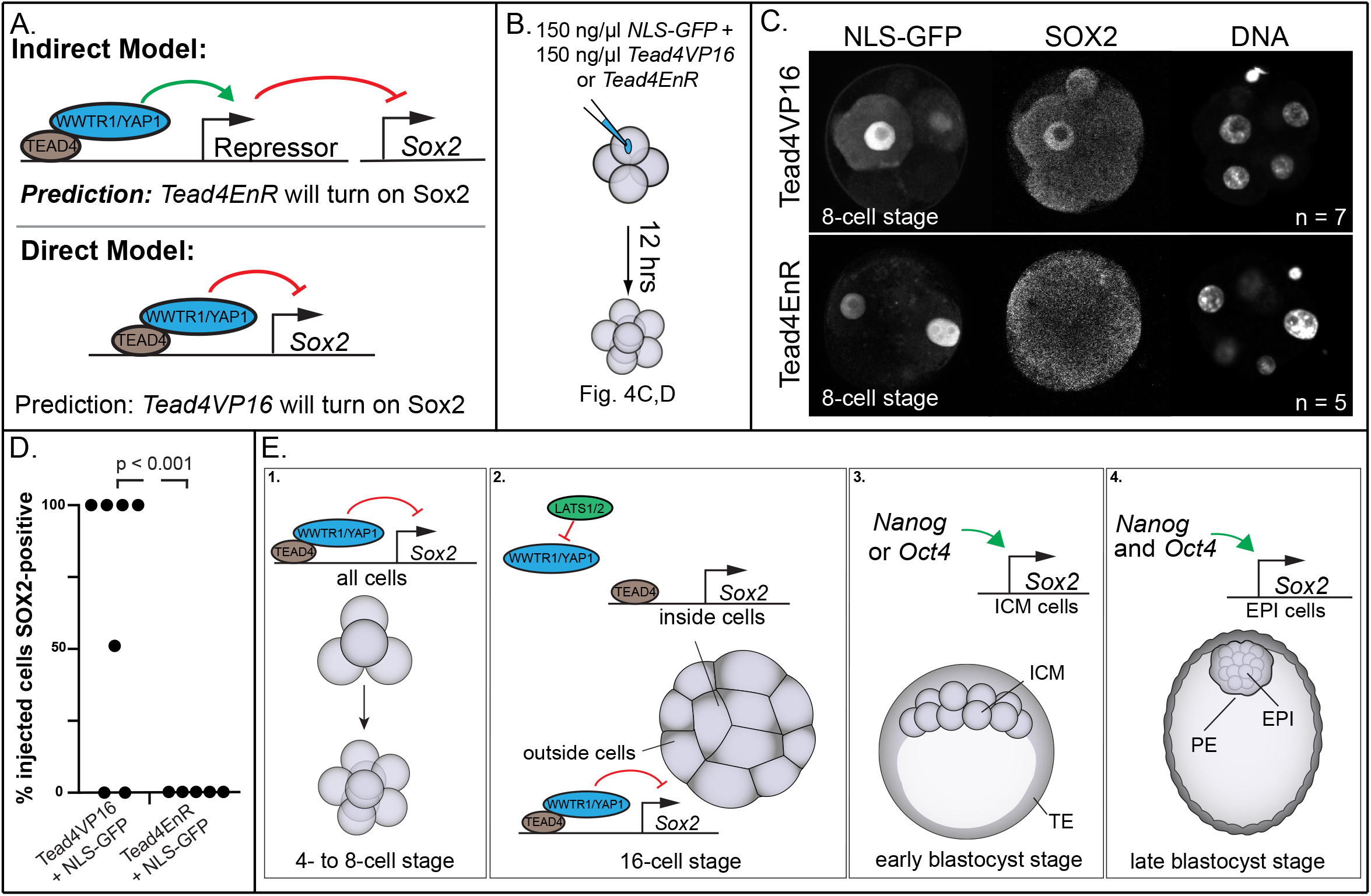
TEAD4/WWTR1/YAP1 repress *Sox2* expression through a direct mechanism. (A) Models for indirect and direct repression of *Sox2* by TEAD4/WWTR1/YAP1 and predicted effect of *Tead4EnR and Tead4VP16* on *Sox2* expression. (B) Experimental approach: a single blastomere of each 4-cell embryo was injected with 150 ng/μl *NLS-GFP* mRNA and either 150 ng/μl *Tead4VP16* or *Tead4EnR* mRNA, and then cultured to the 8-cell stage. (C) GFP and SOX2 immunostaining in embryos injected with *Tead4VP16* or *Tead4EnR*. (D) Quantification of the percentage of NLS-GFP-positive, SOX2-positive cells per embryo (dots) injected with *Tead4VP16* and *Tead4EnR*. p = student’s t-test, n = number of embryos examined. (E) Model for regulation of *Sox2* at indicated developmental stages. ICM = inner cell mass, TE = trophectoderm, EPI = epiblast, PE = primitive endoderm.

To test these models, we employed variants of *Tead4* in which the WWTR1/YAP1 interaction domain has been replaced with either the transcriptional activator VP16 (*Tead4VP16*) or the transcriptional repressor engrailed (*Tead4EnR*). These variants have previously been used in preimplantation embryos to provide evidence that TEAD4/WWTR1/YAP1 promote *Cdx2* expression through a direct mechanism (Nishioka et al., 2009). We reasoned that if TEAD4/WWTR1/YAP1 represses *Sox2* indirectly, then overexpression of *Tead4EnR* would induce *Sox2* expression. Alternatively, if TEAD4/WWTR1/YAP1 represses *Sox2* directly, then *Tead4VP16* would induce *Sox2* expression. We injected mRNA encoding *Tead4VP16* or *Tead4EnR* into a single blastomere of 4-cell stage embryos and tracked progeny of the injected blastomere at the 8-cell stage with co-injection of *GFP* (Fig. 4B). We observed that overexpression of *Tead4VP16*, but not *Tead4EnR* induced SOX2 (Fig. 4C,D). These observations are consistent with the direct repression of *Sox2* by TEAD4/WWTR1/YAP1.

This study highlights distinct phases of *Sox2* regulation occurring during the establishment of pluripotency in mouse development. As early as the 4-cell stage, blastomeres are competent to express *Sox2*, but this is overridden by TEAD/WWTR1/YAP1 (Fig. 4E, box 1). Initiation of *Sox2* expression is independent of *Nanog* and *Oct4* and does not require cues associated with inside cell position or developmental time. Instead, LATS1/2 activity in inside cells relieves repression of TEAD4/WWTR1/YAP1 on *Sox2* (Fig. 4E, box 2). After blastocyst formation, NANOG and OCT4 work together ensure that *Sox2* expression is maintained (Fig 4E, box 3). Finally, as the embryo approaches implantation, *Nanog* and *Oct4* become individually required to sustain *Sox2* expression (Fig. 4E, box 4).

## Materials and Methods

### Mouse strains

Animal care and husbandry was performed in accordance with the guidelines established by the Institutional Animal Care and Use Committee at Michigan State University. Wild type embryos were generated by mating CD-1 mice (Charles River). Female mice used in this study were between six weeks and six months of age and males were used from eight weeks to nine months. Alleles and transgenes used in this study were maintained on a CD-1 background and include: *Nanog^tm1.1Hoch^* (Maherali et al., 2007), *Pou5f1^tm1Scho^* (Kehler et al., 2004), *Tead4^tm1Bnno^* (Yagi et al., 2007), *Yap1^tm1.1Eno^* (Xin et al., 2011), *Wwtr1^tm1.1Eno^* (Xin et al., 2013), *Tg(Zp3-cre)93Knw* (De Vries et al., 2004). Conditional, floxed alleles were recombined to generate null alleles by breeding mice carrying conditional alleles to *Alpl^tm(cre)Nagy^* (Lomelí et al., 2000) mice.

### Embryo collection and culture

Embryos were collected from naturally mated mice by flushing dissected oviducts or uteri with M2 medium (Millipore-Sigma). All embryos were cultured in 5% CO_2_ at 37°C under ES cell grade mineral oil (Millipore-Sigma). Prior to embryo culture, KSOM medium (Millipore-Sigma) was equilibrated overnight in the embryo incubator.

### Embryo microinjection

cDNAs encoding *Lats2, Tead4VP16*, and *Tead4EnR* (Nishioka et al., 2009) cloned into the pcDNA3.1 poly(A)83 plasmid (Yamagata et al., 2005) were linearized, and then used as a template to generate mRNAs for injection by the mMessage mMachine T7 transcription kit (Invitrogen). *NLS-GFP* mRNA was synthesized from linearized *NLS-GFP* plasmid (Ariotti et al., 2015) using the mMessage mMachine Sp6 transcription kit (Invitrogen). Prior to injection, synthesized mRNAs were cleaned and concentrated using the MEGAclear Transcription Clean-up Kit (Invitrogen). *Lats2* and *NLS-GFP* mRNAs were injected into both blastomeres of 2-cell stage embryos at a concentration of 500 ng/μl. *Tead4VP16* or *Tead4EnR* mRNAs were injected into a single blastomere of 4-cell stage embryos at a concentration of or 150 ng/μl each. *NLS-GFP* mRNA was included in 4-cell stage injections at a concentration of 150 ng/μl to trace the progeny of the injected blastomere. All mRNAs were diluted in 10 mM Tris-HCl (pH 7.4), 0.1 mM EDTA. Injections were performed using a Harvard Apparatus PL-100A microinjector.

### Immunofluorescence and Confocal Microscopy

Embryos were fixed in 4% formaldehyde (Polysciences) for 10 minutes, permeabilized with 0.5% Triton X-100 (Millipore-Sigma) for 30 minutes and blocked with 10% FBS, 0.1% Triton X-100 for at least 1 hour at room temperature or longer at 4°C. Primary antibody incubation was performed at 4°C overnight using the following antibodies: goat anti-SOX2 (Neuromics, GT15098, 1:2000), rabbit anti-NANOG (Reprocell, RCAB002P-F, 1:400) mouse anti-Tead4 (Abcam, ab58310, 1:1000), mouse anti-YAP (Santa Cruz, sc101199, 1:200), and rat anti-ECAD (Millipore-Sigma, U3254, 1:500). Anti-SOX2, anti-TEAD4 and anti-YAP antibodies were validated by the absence of positive staining on embryos homozygous for null alleles encoding antibody target Nuclei were labelled with either DRAQ5 (Cell Signaling Technology) or DAPI (Millipore-Sigma). Antibodies raised against IgG and coupled to Dylight 488, Cy3 or Alexa Fluor 647 (Jackson ImmunoResearch) were used to detect primary antibodies. Embryos were imaged on an Olympus FluoView FV1000 Confocal Laser Scanning Microscope using a 20x UPlanFLN objected (0.5 NA) and 5x digital zoom. Each embryo was imaged in entirety using 5 μm optical section thickness.

### Image Analysis

Z-stacks obtained from confocal microscopy were analyzed using ImageJ (Schneider et al., 2012). Each nucleus was identified by DNA stain and then scored for the presence or absence of SOX2. In Fig. 1A and B, cells were classified as inside or outside on the basis of ECAD localization. For analysis of *Nanog;Oct4* null embryos in Fig. 1C,D and Fig. S1D, SOX2 staining intensity was categorized as intense or weak. Intense SOX2 staining was defined as the level observed in non-mutant embryos, which was uniform among inside cells. In Fig. 1, S1, 2, and S2, embryo genotypes were not known prior to analysis. In Fig. 3 and 4 embryos were grouped according to injection performed, and therefore the researcher was not blind to embryo treatment.

### Embryo Genotyping

For embryos at the 8-cell stage or older, DNA was extracted from fixed embryos after imaging using the Extract-N-Amp kit (Millipore-Sigma) in a total volume of 10 μl. For embryos at the 4-cell stage, DNA was extracted from fixed embryos in a total volume of 5 μl. 1 μl of extracted DNA was used as template, with allele specific primers (Table S1).

## Acknowledgements

We thank members of the Ralston Lab for thoughtful discussion. This work was supported by National Institutes of Health (R01 GM104009 and R35 GM131759 to A.R. and T32HD087166 to J.W.).

